# Neural Speech Tracking Across the First 62 Days of Life: Effects of Language Background and Cognitive Maturation

**DOI:** 10.64898/2026.08.04.742719

**Authors:** Martina Dvořáková, Josef Urbanec, Jan Kremláček, Kateřina Chládková

**Author notes:** Correspondence should be addressed to MD.

## Abstract

Newborns recognise familiar language patterns experienced in utero and discriminate them from unfamiliar ones. It is as yet unclear which neural mechanisms are responsible for such early language-specific behavior. Here we focus on neural speech tracking and test whether it shows language-specific attunement in newborns and one-to-two month olds. Infants listened to infant-directed stories in their native language (Czech) and an unfamiliar language (Russian) while their EEG was recorded. First, we examined whether neural speech tracking differed between infants predominantly exposed to Czech and infants exposed primarily to other languages. Second, we assessed developmental changes by combining these data with a previously collected cohort of newborns, yielding a continuous sample spanning 1–62 days of age. Neural speech tracking was assessed in stimulus-derived delta and theta frequency bands in terms of oscillatory power and accuracy of EEG-to-speech envelope reconstruction using the backward multivariate temporal response functions.

Czech-exposed infants had stronger cortical tracking of Czech speech than infants exposed to other languages. Within the Czech-exposed group, native Czech elicited stronger neural tracking than unfamiliar Russian, particularly in the delta band, indicating early language-specific tuning to the prosodic-word structure. Across the combined Czech-exposed sample, this native-language advantage gradually decreased with age, suggesting that neural speech tracking of the slow rhythms undergoes rapid reorganization during the first two months of life. These findings demonstrate that language-specific neural speech tracking is detectable from the earliest weeks of life and is jointly shaped by early, perinatal language experience as well as cognitive maturation.

## 1. Introduction

Humans acquire spoken language with remarkable ease, and converging evidence suggests that the brain is already prepared to process speech at birth (Vouloumanos & Werker, 2007). This early specialization is reflected in newborns’ neural responses, which distinguish natural speech from acoustically matched non-speech sounds and degraded speech signals such as backward or spectrally rotated speech (Chládková et al., 2021; Peña et al., 2003; Sato et al., 2012; May et al., 2018). Beyond this general sensitivity to speech, newborns also show stronger neural responses to the language they were exposed to before birth than to unfamiliar languages (Mehler et al., 1988; Moon et al., 1993; Nazzi et al., 1998; Sato et al., 2012; May et al., 2018). Together, these findings suggest that the infant brain is already selectively sensitive to both speech itself and the specific characteristics of the prenatal linguistic environment.

One explanation for this early language specificity lies in the nature of prenatal auditory experience. During gestation, maternal tissues and amniotic fluid act as a low-pass filter, attenuating higher-frequency acoustic information while preserving temporal cues and low-frequency spectral information below approximately 1 kHz (Richards et al., 1992; Gerhardt & Abrams, 2000; Eggermont & Moore, 2011). Consequently, the fetus is primarily exposed to the rhythm and melody of the surrounding speech. Consistent with this, newborns’ language discrimination is thought to rely predominantly on prosodic information such as rhythm and intonation rather than on segmental detail (Ramus, 2002; Byers-Heinlein et al., 2010; Molnar et al., 2014), although recent work, including computational modelling, suggests that prenatal learning may also extend to at least some vowel categories (Moon et al., 2013; Chládková, Nudga, & Boersma, 2020). Prenatal auditory experience therefore provides a plausible mechanism through which language-specific neural responses can emerge before birth.

Behavioral and neuroimaging studies provide converging evidence for this early language specificity. At the behavioral level, newborns reliably prefer their native language over an unfamiliar one (e.g., Mehler et al., 1988; Bosch & Sebastián-Gallés, 1997; Nazzi et al., 2000; Byers-Heinlein et al., 2010). Likewise, neuroimaging studies demonstrate differential cortical responses to native and unfamiliar speech. For example, hemodynamic responses are stronger or more lateralized for native-language speech presented in its natural forward form than in its backward version (Sato et al., 2012; Vannasing et al., 2016; May et al., 2018), and for native-language speech carrying well-formed rather than foreign or ill-formed prosodic patterns (Martinez-Alvarez et al., 2023; Dvořáková et al., 2006). Together, these findings indicate that language-specific neural processing is already established at birth.

Although these studies demonstrate language-specific neural responses, hemodynamic measures provide only limited information about how the brain processes the rapidly unfolding temporal structure of speech. Neural speech tracking offers a more direct approach by quantifying the extent to which ongoing neural activity aligns with the incoming speech signal. This method captures sensitivity to multiple temporal scales of speech, with slower fluctuations in the delta band (<4 Hz) reflecting phrasal and stress-related structure, and faster modulations in the theta band (approximately 4–8 Hz) corresponding primarily to syllabic organization. By synchronizing with these temporal regularities, neural activity can support the segmentation of continuous speech into meaningful linguistic units such as syllables, words, and phrases (Ding & Simon, 2014; Goswami, 2018; Tune & Obleser, 2022). Consequently, neural speech tracking is thought to facilitate speech processing by supporting the segmentation and identification of linguistic structure (Doelling et al., 2014; Henry & Obleser, 2012; Peelle et al., 2013; Meyer, 2018). As such, it provides a direct and temporally sensitive measure of online speech processing.

Findings from neural speech tracking studies in early infancy remain mixed with respect to language specificity. Some studies suggest that neural tracking is already tuned to the native language. For example, Dvořáková et al. (under review) reported stronger tracking of native Czech than unfamiliar Russian speech in newborns, particularly for infant-directed speech in the delta band. In contrast, other studies have reported little evidence for language-specific neural tracking at birth. Ortiz-Barajas et al. (2021) observed comparable tracking of native and non-native languages in newborns, with language-specific differences emerging only by six months of age. Florea et al. (2024), meanwhile, reported stronger responses to unfamiliar nursery rhymes, suggesting an early bias toward novelty, while also finding enhanced tracking of natural compared with low-pass-filtered versions of familiar rhymes. Together, these findings indicate that although newborns are clearly sensitive to speech rhythms, the extent and nature of language-specific neural speech tracking remain unresolved.

One possible explanation for these inconsistent findings is that newborns differ substantially in their early language experience. To determine whether prior language exposure shapes neural speech tracking, one would ideally compare infants with and without experience of a target language, or infants differing in the amount of exposure they receive. We adopted the latter approach by comparing infants predominantly exposed to Czech with infants growing up in multilingual environments in which Czech constituted only a limited proportion of their early language input. This design allows us to examine whether variation in prenatal and early postnatal language experience influences neural speech tracking at the timescales of prosodic words and syllables.

Neural speech processing is constrained not only by exposure to language, but also by brain maturation. The first weeks after birth are characterized by rapid developmental changes in cognitive capabilities and alertness states (Guyer et al., 2015; Barbeau & Weiss, 2017), which may influence speech processing. The infant brain is not fully mature at birth, and maturational processes are reflected in its electrophysiology (Schaworonkow & Voytek, 2021; Vanhatalo & Kaila, 2006). While slow oscillatory activity in the delta and theta range is already present in auditory and language-related regions prenatally (Arichi et al., 2017; Chipaux et al., 2013; Routier et al., 2017; Vecchierini et al., 2007), faster oscillatory rhythms emerge more gradually over the course of infancy (Le Van Quyen et al., 2006). Consequently, the newborn brain may be particularly well suited to processing speech information unfolding over relatively long temporal windows, such as rhythm, stress patterns, and prosodic structure, while processing rapidly changing segmental information may remain comparatively immature. In line with this, previous work suggests that the development of neural speech tracking follows a similar slow-to-fast rhythm alignment pattern. For example, Attaheri et al. (2022) showed that infants’ tracking of nursery rhymes in the delta band was stronger at 4 months of age than at 7 and 11 months, whereas theta-band tracking remained relatively stable across ages. Delta band tracking seems to play a primary role also in native-language sensitivity in newborns (Dvořáková et al., under review). Studies are lacking that would assess the development in neural speech tracking in the early period between birth and 4 months, during which the infant brain undergoes major development. This period is characterized by rapid neurobiological and behavioral change, including extensive synaptogenesis, myelination, accelerated brain growth (Johnson, 2001; Holland et al., 2014), and rapid developments in speech perception such as perceptual narrowing and increasing sensitivity to subtle language-specific contrasts (Tsuji & Cristia, 2014). Testing how neural speech tracking develops across the first 62 days of life is a second aim of this study.

The present study aims to contribute to developmental neurolinguistics by examining how (1) early language exposure and (2) age shape neural speech tracking during the first two months of life. To address the role of early language experience, we compared infants from monolingual Czech-speaking environments with infants raised in multilingual environments in which Czech constituted only a limited proportion of their early language input. Neural speech tracking was measured using EEG while infants listened to a children’s story presented in Czech and Russian (an unfamiliar language). This design allowed us to determine whether variation in prenatal and early postnatal language exposure modulates neural tracking at the timescales of prosodic words (delta band) and syllables (theta band). We predicted that infants predominantly exposed to Czech would show stronger neural tracking of Czech speech than infants with more limited Czech exposure, reflecting early language-specific tuning. To investigate early developmental change, we examined neural speech tracking across the extended neonatal period (1–62 days). To achieve this, we combined the present sample of infants tested between 3 and 62 days of age with a previously collected cohort of newborns aged 1–6 days who listened to identical stimuli using the same EEG acquisition and analysis procedures. This combined sample enabled us to characterize developmental changes in neural speech tracking continuously across the first two months after birth. Based on previous reports of decreasing delta-band tracking between 4, 7, and 11 months of age (Attaheri et al., 2022), we hypothesized that delta-band tracking would likewise decrease across the neonatal period, although it remains unknown whether such developmental changes are already detectable within the first weeks of life. Together, these complementary analyses allowed us to examine how early language experience and rapid postnatal brain maturation jointly contribute to the emergence of language-specific neural speech tracking.

## 2. Materials and Methods

### 2.1 Stimuli

The auditory materials consisted of naturally spoken Czech and Russian sentences produced in infant-directed speech (IDS) by a native female speaker of each language. The sentences were based on the Czech children’s story *Hrnečku vař!* (a variant of the Brothers Grimm tale *Sweet Porridge*) and were translated into Russian by a balanced bilingual speaker of Russian (L1) and Czech (L2). The story was divided into 70 sentences and presented three times throughout the experiment. The original sentence sequence was maintained to preserve the narrative structure and natural prosodic continuity of the story.

To approximate natural listening conditions and minimize artificial temporal predictability, pauses between consecutive sentences were brief and varied randomly between 350 and 400 ms in 25-ms increments. Infants completed three experimental blocks: a control block containing white noise lasting approximately 3 minutes, followed by two speech blocks of approximately 10 minutes each, one in Czech and one in Russian. Presentation order of the two language blocks was counterbalanced across participants. The experimental design is shown in Figure 1 below, and the acoustic properties of the stimuli are summarized in Table 1 below.

**Figure 1.**
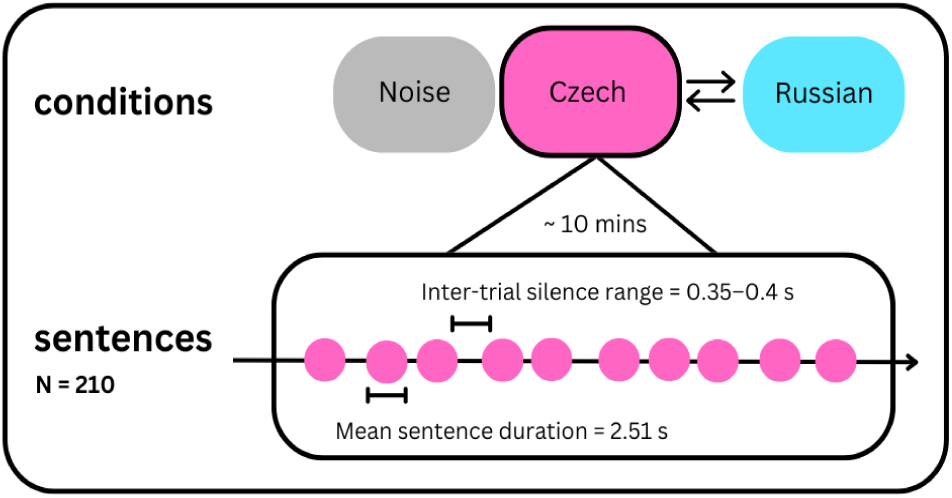
Experiment design: the Noise condition was always presented as the first block, followed by blocks of Czech (familiar language) and Russian (unfamiliar language), counterbalanced between infants.

**Table 1.** Duration and F0 of the stimuli, means and 95% confidence intervals.

|  | **Czech**<br>**Speech** | **Russian**<br>**Speech** |
| --- | --- | --- |
| **Sentence duration**<br>**(s)** | 2.51<br>[2.42, 2.59] | 2.68<br>[2.57, 2.78] |
| **Syllable rate (n**<br>**sylls/s)** | 4.94<br>[4.8, 5.07] | 4.65<br>[4.53, 4.77] |
| **Word rate (n**<br>**words/s)** | 1.91<br>[1.84, 1.99] | 1.6<br>[1.55, 1.65] |
| **F0 mean (Hz)** | 200.08<br>[194.75,<br>205.40] | 183.85<br>[181.08,<br>186.62] |
| **F0 range (Hz)** | 165.13<br>[151.14,<br>179.12] | 170.93<br>[155.38,<br>186.49] |

### 2.2 Participants

The infants recruited for this study were 57 healthy one-month olds, born into monolingual or multilingual families in the Czech Republic^1^ with Czech as the dominant society language. Sixteen infants were excluded due to hardware malfunction (*n* = 1), and due to providing insufficient data (*n* = 15; see “EEG preprocessing and Power” section below). The final sample comprised 41 infants (22 males, 19 females) whose mean age was 32.34 days, ranging from 3 to 62 days; all were full term, born at GA > 36 weeks via vaginal delivery or uncomplicated cesarean section.

Infants were considered at low risk for developmental language or speech disorders based on the absence of such conditions in their parents and siblings. The sample consisted of 30 infants in the Czech-exposed Group, whose mothers were native Czech speakers and reported speaking Czech more than 60% of time (on average 90.2 % of spoken Czech a day during third trimester and until the testing session based on mothers’ self-evaluation during pregnancy and the testing session). There were another 11 infants in the Other-exposed Group, whose mothers spoke more than 60% of time in one or more languages other than Czech (namely, English [5]; English and Slovak [4]; Slovak [1]; Russian [1]).

An additional group of one-to-tree day old newborns was included in the present analyses of the age effect. This data represents a subset of the sample of our earlier study on infant-vs adult-directed speech processing in newborns (Dvořáková et al., under review; only the infants from the IDS condition of the previous study are analyzed here). These are 27 newborns (13 males, 14 females; mean age = 2.78 days, range = 1–6 days; mean gestational age at birth = 39.42 weeks, range = 36–41 weeks; mean birth weight = 3190 g, range = 2480–4380 g). All mothers were native Czech speakers and functional monolinguals. Across the combined Czech-exposed sample (the Czech-exposed Group and the newborn group), infants ranged in age from 1 to 62 days (mean = 20.3 days, median = 21 days; interquartile range = 3–33 days). Table 2 gives an overview of demographic and data quality measures of the three infant samples assessed in the present study.

**Table 2.** Demographic characteristics and data quality measures for the three participant groups included in the study.

|  | Final N<br>Included<br>in the<br>Analyses | M Age<br>[days] | Sex<br>Assigned<br>at Birth | Included<br>Trials per<br>Condition | % daily<br>exposure to<br>Czech | N of<br>Excluded<br>Infants<br>due to<br>Artifacts |
| --- | --- | --- | --- | --- | --- | --- |
| Czech-exposed<br>one-month olds | 30 | 33.63<br>(SD=12) | 15 M, 15 F | 160 (SD=39) | 90.2 (SD=11.7) | 8 |
| Other-exposed<br>one-month olds | 11 | 28.82<br>(SD=15.85) | 7 M, 4 F | 159 (SD=45) | 16 (SD=17.6) | 7 |
| Czech-learning newborns | 25 | 2.84 (SD=1.58) | 12 M, 13 F | 154 (SD=40) | 98.8 (SD=4.4) | 2 |

### 2.3 Design Considerations and Sample Size

This study assesses neural speech tracking data of a total of 66 infants, 41 one-month olds (recruited for this particular study, further split in two groups differing in language background) and 25 newborns (a planned comparison addressing the age affect; a subset of data collected in an earlier study on neural tracking of IDS vs ADS). The present study employed a repeated-measures EEG design in which each infant contributed neural speech-tracking estimates for two languages (Czech and Russian) and two frequency bands (delta and theta). This within-subject design reduces between-participant variability and increases sensitivity to detect experimental effects relative to a purely between-subject comparison. In addition, each infant contributed approximately 160 artifact-free trials per condition, providing reliable within-participant estimates of neural speech tracking.

### 2.4 Procedure

The one-month old infants were tested in a linguistics babylab at Charles University in Prague^2^. During testing, an infant was either asleep or awake, with the states of sleep and wakefulness changing during testing. They were held by their mother in a comfortable supine position. Stimuli were presented through Kurzweil loudspeakers placed to the left and right in front of the infant; the sound level was at 67 dB SPL as measured at the position of the infant’s head.

The younger, newborn cohort was recruited and tested at the maternity ward of Havlíčkův Brod Hospital (these infants are a subset of the sample analysed in a previous study on neural tracking of infant-vs adult-directed speech, Dvořáková et al., under review). Newborns were recorded while asleep in their hospital cots; a mother was always present, seated in a chair in proximity to the infant. Auditory stimulation was presented binaurally through ER-3C insert earphones (Etymotic Research, Inc.) equipped with disposable Flexicouplers (Natus Europe GmbH), with the sound level set to 67 dB SPL.

Prior to participation, mothers provided written informed consent. The study protocol was approved by the Ethics Committee of the Institute of Psychology of the Czech Academy of Sciences (PSU-925/Brno/2022) and complied with the ethical standards outlined in the Declaration of Helsinki (2008).

### 2.5 EEG preprocessing

EEG preprocessing and analysis followed the procedures described in Dvořáková et al. (under review). Continuous EEG was acquired from six Ag/AgCl scalp electrodes positioned at F3, Fz, F4, C3, Cz, and C4. Signals were referenced to a facial electrode placed on the cheek in older infants and on the nose in newborns. Recordings were obtained using a DEYMED Diagnostic amplifier (DEYMED Diagnostic Ltd., Czech Republic) with an acquisition bandwidth of 0.3–100 Hz.

All analyses were conducted in MATLAB R2024b (The MathWorks, Inc., Natick, MA) using custom-written scripts together with functions from EEGLAB (Delorme & Makeig, 2004) and the mTRF Toolbox (Crosse et al., 2016). The EEG data were downsampled to 200 Hz and filtered between 0.05 and 30 Hz. Electrodes showing absent or negligible signal variability (standard deviation < 10⁻⁶ V) were classified as flat channels and reconstructed using spherical interpolation.

Frequency bands of interest were defined based on the temporal characteristics of the speech stimuli. Specifically, speech rates corresponding to word-level and syllable-level structure were used to derive delta- and theta-band ranges, respectively (Table 1). To accommodate variability across the Czech and Russian materials, the stimulus-derived ranges were extended by ±1 Hz and combined across languages by taking the lowest and highest resulting boundaries. This procedure yielded a common delta band of 0.70–2.99 Hz and a common theta band of 3.53–6.07 Hz, which were subsequently applied to all analyses.

### 2.6 Power

For the time–frequency analyses, the continuous EEG data were divided into epochs extending from 300 ms before sentence onset to 3700 ms after onset. Each epoch therefore encompassed the neural response to an individual sentence together with short periods of preceding and following silence. A 4-s epoch length was chosen to ensure adequate representation of low-frequency oscillatory activity in subsequent spectral analyses. Epochs containing amplitudes exceeding ±350 µV were discarded. Participants had to contribute more than 30% of artifact-free epochs; on average, 160 epochs (SD = 39) per condition were retained for infants in the Czech-learning one-month-old group, 159 epochs (SD = 45) for infants in the Other-language one-month-old group, and 154 (SD=40) for infants in the Czech-learning newborn group.

Time–frequency analyses were carried out in EEGLAB (Delorme & Makeig, 2004) using the pop_newtimef function. Event-related spectral perturbations (ERSPs) were estimated separately for each participant, condition, and electrode using Morlet wavelets. The wavelet width increased progressively from 2 cycles at the lowest analyzed frequency (0.7 Hz) to 8.7 cycles at the highest frequency (10.2 Hz). Spectral estimates were computed across the interval from −300 to 3700 ms and frequencies ranging from 0.7 to 10.2 Hz in 0.1-Hz increments.

Because of the short intervals between sentences no baseline correction was applied. Instead, speech-related changes in oscillatory power were quantified by calculating the difference in power (dB) between the speech and noise conditions, effectively expressing speech-evoked activity relative to background neural activity. Time–frequency estimates were evaluated at 163 valid temporal locations returned by EEGLAB. The number of output locations was constrained by the duration of the wavelet at the lowest analysed frequency. A padding ratio of 4 was used to increase the density of frequency sampling.

### 2.7 multivariate Temporal Response Function (mTRF)

To quantify the extent to which neural activity tracked the temporal dynamics of speech, we employed backward multivariate temporal response functions (mTRFs). Unlike forward encoding approaches, which predict neural responses from the stimulus, the backward reconstruction model estimates the speech envelope from the EEG signal. Because information from all electrodes is combined into a single decoder, backward mTRFs generally provide higher reconstruction accuracy and improved sensitivity to stimulus–brain coupling (Di Liberto et al., 2015; Crosse et al., 2016; Kalashnikova et al., 2018; Jessen et al., 2019; Attaheri et al., 2022).

Analyses were conducted separately for the delta and theta frequency ranges using stimulus-specific frequency bands applied to both EEG recordings and speech envelopes. Channels whose activity exceeded three standard deviations from the participant-specific mean were classified as outliers and replaced using spherical spline interpolation. The EEG recordings were then divided into segments consisting of nine consecutive sentences. This procedure yielded average segment lengths of 26.14 s for Czech stimuli and 27.62 s for Russian stimuli, with 23 segments for each condition. Speech envelopes were derived using the Hilbert transform.

Prior to envelope extraction, the audio signal was band-pass filtered using a third-order Butterworth high-pass filter and a fifth-order Butterworth low-pass filter. The resulting envelopes were downsampled to 200 Hz to match the sampling frequency of the EEG data. EEG and envelope signals were subsequently aligned and truncated to equal duration. Following previous infant mTRF studies (Di Liberto et al., 2015; Attaheri et al., 2022; Dvořáková et al., under review), the decoder was trained using time lags ranging from 0 to 250 ms. Both neural and acoustic signals were standardized using z-score normalization before model fitting.

To reduce overfitting, model estimation relied on ridge regression. For each participant, condition, and frequency band, a leave-one-out cross-validation procedure was applied across stimulus segments. In each iteration, one segment was reserved for testing while the remaining segments were used to train the decoder. Model performance was evaluated across a logarithmically spaced range of regularization parameters (λ = 10⁻³–10⁸). For every λ value, the reconstructed envelope was correlated with the original speech envelope using Pearson’s correlation coefficient. The λ producing the highest average reconstruction accuracy across folds was selected as optimal, and its corresponding correlation coefficient served as the measure of neural speech tracking used in subsequent analyses.

### 2.8 Statistical analyses

Data were analyzed using linear mixed-effects (LME) models in R with the packages *lme4* and *lmerTest* (Bates et al., 2015; Kuznetsova et al., 2017; R Core Team, 2016). Estimated means and confidence intervals were obtained using the package *ggeffects* (Lüdecke, 2018) and visualized with *ggplot2* (Wickham, 2016).

One set of models was run to test the effects of Language Background, another set of models was done to assess the effect of Age. Within each set of analyses, one model was fitted for power and one was fitted for mTRF. The analyses of Language background included the data of the Czech-exposed and Other-exposed one-month olds. The fixed effects were Band, Condition, Background, Block Order, and all their interactions. The factors Band, Condition and Background were treatment-coded with Delta (vs Theta), Czech stimuli (vs Russian stimuli), and Czech-language background (vs Other-language background) as reference levels, respectively, and Block Order was coded orthogonally as first vs. second block. The random-effects structure included by-participant random intercepts and by-participant random slopes for Condition and Band for mTRF and by-participant random slopes for Condition and Band and their interaction for power.

The analyses of Age were done on the data of Czech-learning one-month olds and Czech-learning newborns. The fixed effects were Band, Condition, Age Group, Block Order, and all their interactions. The factors Band, Condition were treatment-coded with Delta and Czech stimuli as reference levels, respectively, Block Order was coded orthogonally as first vs. second block and Age was a continuous numeric factor and z-scored. The random-effects structure included by-participant random intercepts, and for mTRF model by-participant random slopes for Condition, and for the power model by-participant random slopes for Condition and Band and their interaction.

## 3. Results

### 3.1 Language Background

Fixed-effect summaries of the models for Language Background are reported in Table 3. Figure 2 shows the grand-averaged measured (and denoised) oscillatory power, and Figure 3 visualises the model-estimated means and 95% confidence intervals for power and mTRF.

**Figure 2.**
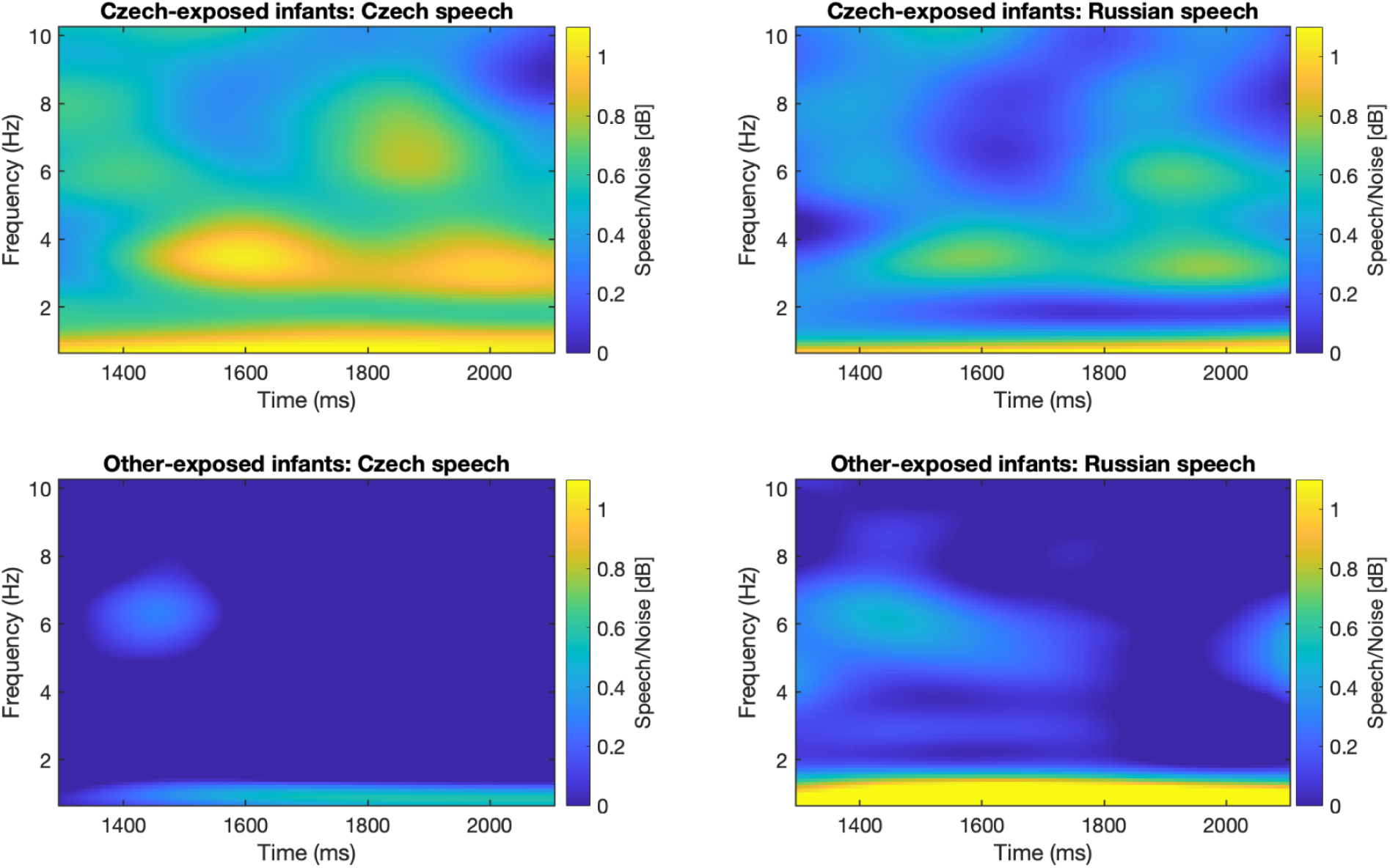
Grand-average ratio between Speech and Noise EEG Power across conditions and backgrounds. Time–frequency representations of grand-average neural responses to naturally produced familiar-language/Czech and unfamiliar-language/Russian sentences in Czech-exposed and Other-exposed one-month old infants. Panels show the speech-to-noise ratio [differences in dB] across time and frequency for the four experimental conditions. Warmer colours indicate stronger positive speech-related responses, whereas cooler colours indicate weaker values.

**Figure 3.**
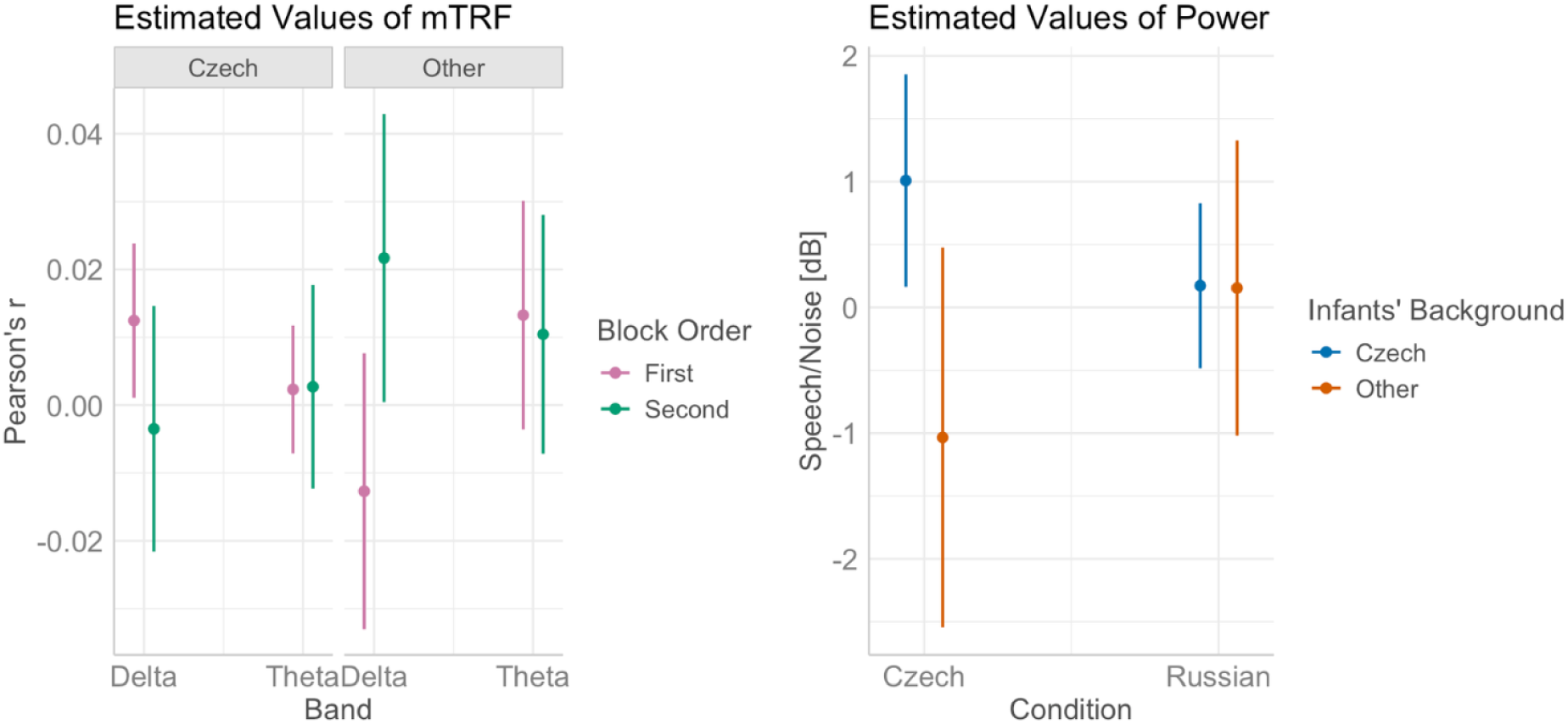
Left: Estimated marginal means and 95% confidence intervals for backward mTRF, showing the model-predicted Pearson’s r values per Band, Background, and Block Order. **Right:** Estimated marginal means and 95% confidence intervals for ratio between Speech and Noise EEG Power.

**Table 3.** Fixed-effect summaries for the linear-mixed effect models on mTRF and Power.

| Effect | mTRF |  |  |  |  | Power |  |  |  |  |
| --- | --- | --- | --- | --- | --- | --- | --- | --- | --- | --- |
|  | Est. | SE | df | *t* | *p* | Est. | SE | df | *t* | *p* |
| Intercept (Condition = Czech, Band = Delta, Background = Czech) | 0.004 | 0.006 | 47.600 | 0.779 | .440 | 0.709 | 0.431 | 36.997 | 1.646 | .108 |
| ConditionRussian | 0.005 | 0.007 | 68.910 | 0.662 | .510 | -0.330 | 0.368 | 37.001 | -0.897 | .376 |
| BandTheta | -0.002 | 0.006 | 76.630 | -0.311 | .756 | -0.049 | 0.218 | 36.995 | -0.226 | .823 |
| BackgroundOther | 0.000 | 0.012 | 47.600 | 0.001 | .999 | -1.150 | 0.882 | 36.997 | -1.304 | .200 |
| BlockOrder (First vs Second) | -0.008 | 0.006 | 47.600 | -1.385 | .173 | -0.300 | 0.431 | 36.997 | -0.696 | .491 |
| Condition × Band | -0.006 | 0.008 | 74.000 | -0.746 | .458 | 0.092 | 0.187 | 37.001 | 0.488 | .628 |
| Condition × Background | 0.004 | 0.015 | 68.910 | 0.252 | .802 | 1.564 | 0.754 | 37.001 | 2.074 | .045 |
| Band × Background | 0.009 | 0.013 | 76.630 | 0.718 | .475 | -0.003 | 0.446 | 36.995 | -0.008 | .994 |
| Condition × BlockOrder | 0.015 | 0.008 | 37.020 | 1.772 | .085 | 0.506 | 0.678 | 36.996 | 0.746 | .460 |
| Band × BlockOrder | 0.008 | 0.006 | 76.630 | 1.288 | .202 | 0.308 | 0.218 | 36.995 | 1.414 | .166 |
| Background × BlockOrder | 0.025 | 0.012 | 47.600 | 2.134 | .038 | 0.893 | 0.882 | 36.997 | 1.013 | .318 |
| Condition × Band × Background | -0.009 | 0.017 | 74.000 | -0.519 | .605 | -0.722 | 0.384 | 37.001 | -1.882 | .068 |
| Condition × Band × BlockOrder | -0.017 | 0.010 | 42.830 | -1.758 | .086 | -0.500 | 0.380 | 36.995 | -1.315 | .196 |
| Condition × Background × BlockOrder | -0.027 | 0.017 | 37.020 | -1.606 | .117 | -0.460 | 1.388 | 36.996 | -0.331 | .742 |
| Band × Background ×<br>BlockOrder | -0.027 | 0.013 | 76.630 | -2.061 | .043 | -0.187 | 0.446 | 36.995 | -0.419 | .678 |
| Condition × Band ×<br>Background ×<br>BlockOrder | 0.037 | 0.020 | 42.830 | 1.820 | .076 | -0.345 | 0.779 | 36.995 | -0.443 | .660 |

For Language Background, the Power analysis revealed a significant Condition × Background interaction (β = 1.564, SE = 0.754, *t*(37.00) = 2.07, *p* = .045). Pairwise comparisons, visualized in Figure 3 (right), showed that for the Czech stimuli, Czech-exposed infants had stronger oscillatory power than Other-exposed infants. The mTRF analysis revealed a significant interaction between Background and Block Order (β = 0.025, SE = 0.012, *t*(47.60) = 2.13, *p* = .038); pairwise comparisons of estimated means, visualised in Figure 3 (left), indicated that Other-exposed infants had stronger neural tracking to the condition presented as second. This effect was licensed by a higher-order interaction between Frequency band, Background, and Block Order (β = −0.027, SE = 0.013, *t*(76.58) = −2.06, *p* = .043); comparisons of the estimated means in Figure 3 (left) suggested that the stronger tracking of the condition presented as second in the Other-language exposed infants was specific to the delta band.

### 3.2 Age

Fixed-effect summaries of the models for Age are reported in Table 4. Figure 4 shows the grand-averaged measured (and denoised) oscillatory power, and Figure 5 visualises the model-estimated means and 95% confidence intervals for power and mTRF.

**Figure 4.**
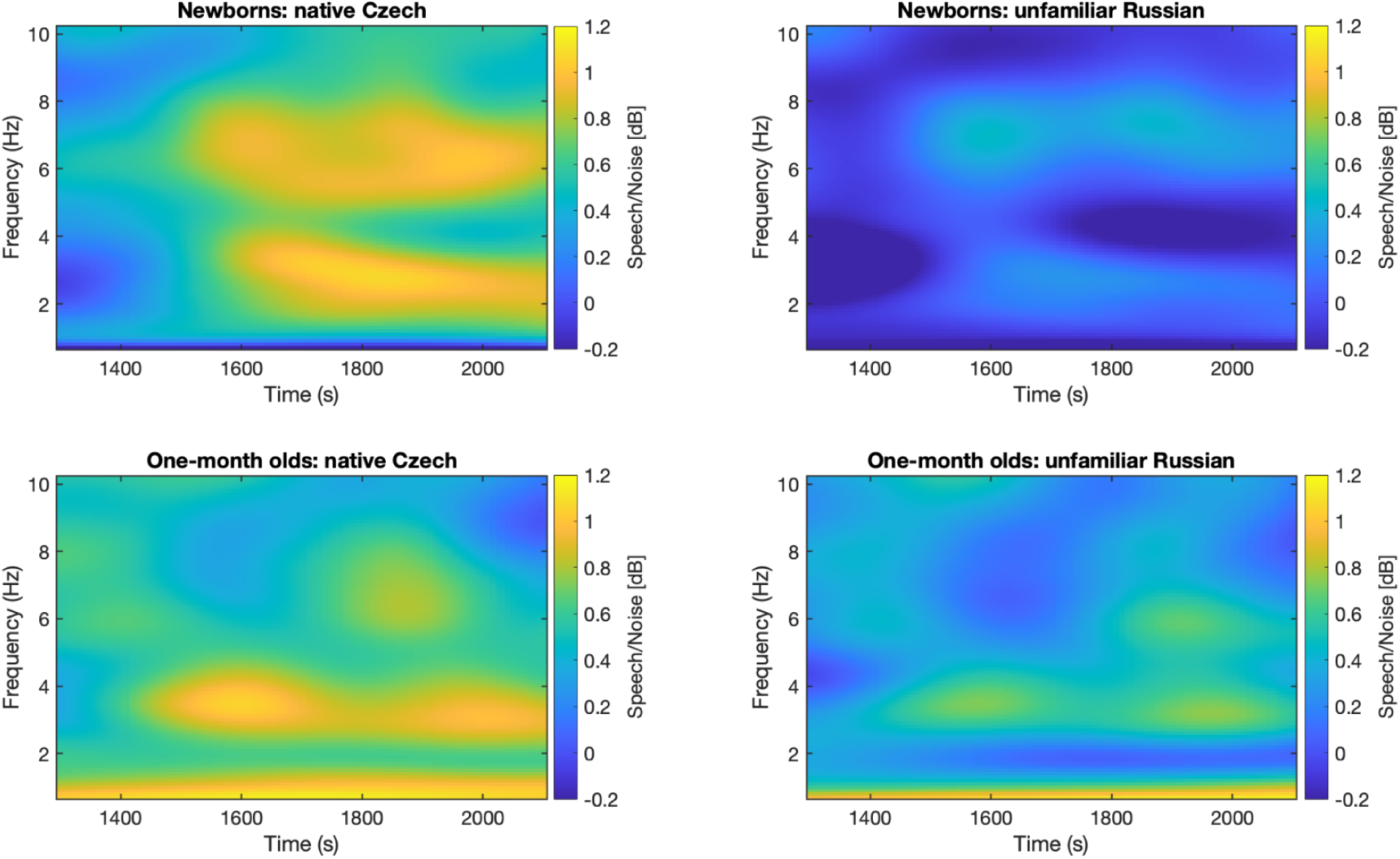
Grand-average ratio between Speech and Noise EEG Power across conditions and backgrounds. Time–frequency representations of grand-average neural responses to naturally produced Czech (CZ) and Russian (RU) sentences in newborns and one-month olds. Warmer colours indicate stronger positive speech-related responses, whereas cooler colours indicate weaker or negative values.

**Figure 5.**
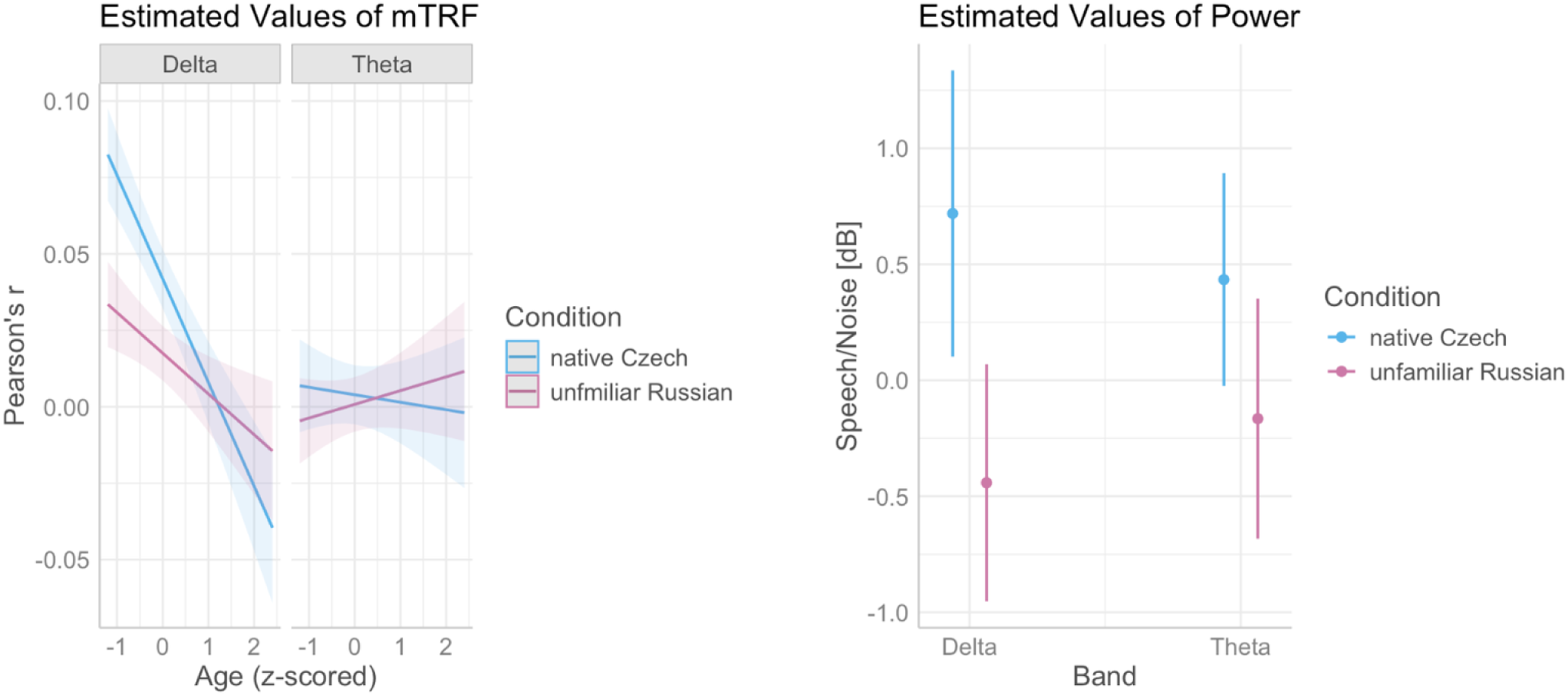
Left: Estimated marginal means and 95% confidence intervals for backward mTRF, showing the model-predicted Pearson’s r values per Condition, Band, and Age. **Right:** Estimated marginal means and 95% confidence intervals for ratio between Speech and Noise EEG Power.

**Table 4.** Fixed-effect summaries for the linear-mixed effect models on mTRF and Power.

|  | mTRF |  |  |  |  | Power |  |  |  |  |
| --- | --- | --- | --- | --- | --- | --- | --- | --- | --- | --- |
| Effect | Est. | SE | df | t | p | Est. | SE | df | t | p |
| (Intercept) | 0.043 | 0.005 | 104.910 | 8.701 | < .001 | 0.699 | 0.315 | 49.188 | 2.220 | .031 |
| Condition (RU vs CZ) | -0.016 | 0.007 | 109.300 | -2.416 | .017 | -0.559 | 0.265 | 41.276 | -2.107 | .041 |
| Band (Theta vs Delta) | -0.040 | 0.006 | 96.091 | -6.578 | < .001 | -0.053 | 0.209 | 48.354 | -0.254 | .800 |
| Age | -0.037 | 0.005 | 108.900 | -7.651 | < .001 | 0.247 | 0.287 | 53.676 | 0.861 | .393 |
| Block order (Second vs First) | 0.001 | 0.005 | 115.500 | 0.182 | .856 | -0.004 | 0.225 | 88.285 | -0.016 | .987 |
| Condition × Band | 0.015 | 0.009 | 96.091 | 1.718 | .089 | 0.022 | 0.144 | 47.418 | 0.154 | .878 |
| Condition × Age | 0.022 | 0.007 | 113.200 | 3.258 | .001 | -0.111 | 0.248 | 46.047 | -0.448 | .656 |
| Band × Age | 0.036 | 0.006 | 96.091 | 6.021 | < .001 | -0.160 | 0.196 | 53.938 | -0.815 | .419 |
| Condition × Block order | 0.008 | 0.007 | 122.600 | 1.217 | .226 | 0.583 | 0.360 | 83.810 | 1.619 | .109 |
| Band × Block order | -0.002 | 0.006 | 96.091 | -0.381 | .704 | 0.225 | 0.173 | 82.841 | 1.301 | .197 |
| Age × Block order | -0.003 | 0.005 | 108.400 | -0.636 | .526 | -0.572 | 0.291 | 52.777 | -1.965 | .055 |
| Condition × Band × Age | -0.020 | 0.009 | 96.091 | -2.341 | .021 | 0.130 | 0.139 | 51.645 | 0.931 | .356 |
| Condition × Band × Block order | -0.006 | 0.009 | 96.091 | -0.730 | .467 | -0.533 | 0.336 | 85.306 | -1.586 | .116 |
| Condition × Age × Block order | 0.001 | 0.006 | 117.000 | 0.176 | .861 | 0.793 | 0.474 | 51.102 | 1.674 | .100 |
| Band × Age × Block order | 0.005 | 0.006 | 96.091 | 0.818 | .415 | 0.147 | 0.198 | 52.877 | 0.743 | .461 |
| Condition × Band × Age × Block order | -0.006 | 0.009 | 96.091 | -0.745 | .458 | -0.166 | 0.384 | 52.994 | -0.433 | .667 |

On the data for Age, the Power analysis revealed a significant intercept (β = 0.699, SE = 0.315, t = 2.220, p = .031), indicating reliable neural tracking in all infants in terms of power in the delta band to Czech speech. A significant main effect of Condition showed stronger tracking to the familiar-language, Czech speech than to the unfamiliar-language, Russian speech in delta band, the reference level (β = −0.559, SE = 0.265, t = −2.107, p = .041).

In the mTRF analysis, there was a significant intercept (β = 0.043, SE = 0.005, t = 8.70, p < .001), again indicating that neural speech tracking in the reference configuration (CZ condition, Delta frequency), averaged across age and block order, was significantly above zero. The model revealed a significant main effect of Condition (β = −0.016, SE = 0.007, t = −2.42, p = .017), showing that Czech speech was tracked more precisely than Russian speech in delta band; significant main effect of Band (β = −0.040, SE = 0.006, t = −6.58, p < .001) indicated better tracking in delta band compared to theta band in the Czech condition (the reference level). A main effect of Age showed a decrease in neural speech tracking as age increased (β = −0.037, SE = 0.005, t = −7.65, p < .001).

These effects were qualified by significant interactions, including a Condition × Age interaction (β = 0.022, SE = 0.007, t = 3.26, p = .001), showing that with increasing age, the difference between responses to Czech and Russian was smaller. A significant Condition × Band × Age interaction further licensed this effect (β = −0.020, SE = 0.009, t = −2.34, p = .021), indicating that the attenuated difference between neural tracking of Czech vs Russian in older infants was specific to the delta band. There was also a significant Band × Age interaction (β = 0.036, SE = 0.006, t = 6.02, p < .001), showing that with increasing age, the difference between delta and theta decreased.

## 4. Discussion

This study investigated neural speech tracking in young infants while they listened to children’s stories in infant-directed native-language (Czech) and unfamiliar-language (Russian) speech. We tested whether cortical speech tracking is modulated by infants’ language background and by their age, comparing, respectively, one-month old infants learning Czech to one-month old infants learning also other languages, and Czech-learning one-month olds to Czech-learning newborns. We found that infants with Czech exposure showed stronger cortical tracking of Czech than infants exposed to other languages. Moreover, within the Czech-exposed group, infants tracked their native-language Czech speech more strongly than they tracked unfamiliar-language Russian speech. Finally, the language-specific advantage for native over unfamiliar speech became smaller with increasing age.

The present study demonstrates that neural speech tracking is shaped by the earliest language experience: one-month old infants who had been exposed to Czech showed enhanced tracking of Czech speech unlike infants with multilingual exposure. This finding extends previous behavioral and neuroimaging evidence by demonstrating that language experience shapes cortical tracking of continuous speech already within the first month of life (Moon et al., 1993; Byers-Heinlein et al., 2010; Sato et al., 2012; Vannasing et al., 2016; May et al., 2018).

Importantly, language-specific tuning was also evident within the Czech-exposed infants who had stronger cortical tracking of their native Czech than of unfamiliar Russian speech in the delta band, corresponding to the temporal rate of prosodic words. This finding indicates that the youngest infants (namely, between 0 and 2 months of age, which was the age range in the present Czech-exposed sample) are not only broadly sensitive to speech but already show selective neural tuning to the prosodic properties of their native language at the level of word rhythm (Ortiz-Barajas et al., 2023; Dvořáková et al., under review). Although previous work has reported mixed results regarding the presence of language-specific effects at birth (Ortiz-Barajas et al., 2021; Florea et al., 2024), our findings support the view that neural speech tracking can capture subtle early differences in how the native language is processed, in contrast to another natural but unfamiliar language.

Although the primary effects of language exposure followed our predictions, an additional and unexpected pattern emerged in the other-language exposure group, where infants showed stronger tracking for the second stimulus block, particularly in the delta band. A possible explanation could lie in the greater variability of the linguistic environment experienced by these infants. Infants in the Other-exposed group typically heard one or more language(s) other than Czech at home while also being exposed to some Czech through the wider community. Such multilingual experience has been associated with attenuated perceptual narrowing and prolonged sensitivity to non-native speech contrasts (e.g., Byers-Heinlein &Fennell, 2014; Graf Estes & Hay, 2015), potentially resulting in greater flexibility and adaptability when processing successive speech streams. This increased adaptability to variable linguistic input may have contributed to the enhanced neural tracking observed in the second stimulus block. While lending itself as an interesting ground for future research, this block effect will not be discussed here in more detail as it did not involve the Language Background by Condition interaction, which is our research-question answering parameter.

The second major finding concerns developmental changes in neural speech tracking. We observed an age-related decrease in the difference between stronger tracking of native-language speech and weaker tracking of unfamiliar-language speech in the delta band. This result is broadly consistent with previous developmental work showing changes in low-frequency speech tracking across infancy. For example, Attaheri et al. (2022) reported that delta-band tracking of infant-directed speech peaks earlier in development and decreases over the first year, whereas theta-band tracking remains relatively stable. Our findings extend this developmental trajectory to a much earlier period, suggesting that cortical speech tracking begins to reorganize already within the first two months after birth.

The attenuation in language-specific delta tracking within the first two months after birth likely reflects a combination of neural maturation, cognitive states, and/or stages of language acquisition. One possible explanation relates to maturational changes in the electrophysiological properties of the infant brain. Delta activity is particularly prominent during fetal life and the early postnatal period (Arichi et al., 2017; Vecchierini et al., 2007). During this stage, neural activity is dominated by slow oscillatory patterns that gradually become reorganized as cortical networks mature (Schaworonkow & Voytek, 2021; Anderson & Perone, 2018). Early neural systems may therefore be especially sensitive to the slow temporal structure of speech, including prosodic and rhythmic information that is abundantly available in the prenatal environment (Granier-Deferre et al., 2011). As maturation progresses, speech processing may become less dominated by these low-frequency mechanisms, reducing the relative advantage of the slowest native-language rhythms observed in the youngest infants. Consequently, the earliest stages of language acquisition may rely disproportionately on slow temporal information, such as rhythm and prosodic structure, with progressively finer-grained linguistic representations becoming available as cortical networks mature (Menn et al., 2023a). The decreasing difference between tracking native Czech and unfamiliar Russian speech should therefore not be interpreted as a loss of language-specific tuning. Rather, it likely reflects a developmental reorganization of the neural mechanisms supporting speech processing, whereby language-specific information is progressively represented through increasingly differentiated neural processes.

Although neural maturation provides one plausible explanation, developmental changes in sleep and behavioral state may also have contributed to the observed age effects. Practically all of the youngest newborns were tested asleep, whereas older infants were more frequently awake, restless, or transitioning between wakefulness states. Newborns spend approximately 70% of their time asleep, compared to roughly 57% by one month of age (Guyer et al., 2015). Moreover, sleep architecture undergoes developmental changes during this period. In newborns, active and quiet sleep occupy approximately equal proportions of sleep time, whereas the proportion of quiet sleep gradually increases with age (Barbeau & Weiss, 2017). Quiet sleep in newborns is characterized by high-amplitude slow-wave activity dominated by delta frequencies, as well as by *tracé alternant*, a discontinuous EEG pattern consisting of bursts of high-amplitude delta and theta activity interspersed with lower-amplitude faster activity (Vanhatalo & Kaila, 2006; Vecchierini et al., 2007). This slow-activity dominance may have amplified delta-band tracking in the youngest infants. Sleep-related physiology therefore likely modulated the magnitude of speech tracking without altering its fundamentally language-specific organization. Future longitudinal studies combining detailed measures of behavioral state, neural maturation, and language development will be necessary to disentangle these mechanisms and determine how early speech-tracking responses are reorganized during the first months of life.

In sum, the present findings suggest that cortical speech tracking provides a sensitive neural marker of the interaction between early language experience and brain maturation. Rather than reflecting fixed neural representations, speech tracking appears to capture a rapidly developing system in which language-specific tuning emerges immediately after birth and is subsequently refined as cortical networks mature.

## 5. Conclusions

Neural speech tracking is shaped by both language experience and age within the first two months of infant development. Infants acquiring Czech showed stronger tracking of their native-language Czech speech than did infants exposed multilingually, primarily to other languages. Czech-exposed infants also tracked more strongly their native language compared to an unfamiliar language. At the same time, the advantage of strongest neural tracking of the native language as opposed to an unfamiliar language decreased across the first two months after birth, probably due to a combined effect of physiological and cognitive factors.

## Acknowledgements

We would like to thank the participating families for their trust and willingness to bring their babies to our babylab within weeks after birth.

## Data and Code Availability Statements

Our stimuli and the custom preprocessing and analysis scripts are available at https://osf.io/sc2p5/overview?view_only=dd3a110630f547fb9f26abaa23a6a272.

## Author Contribution

**Martina Dvořáková**: conceptualization, investigation, formal analysis, funding acquisition, methodology, project administration, resources, visualization, writing – review and editing, writing – original draft. **Josef Urbanec**: investigation, project administration, writing – review and editing, resources. **Jan Kremláček**: data curation, formal analysis, writing – review and editing. **Kateřina Chládková**: conceptualization, formal analysis, funding acquisition, methodology, resources, supervision, visualization, writing – original draft, writing – review and editing.

## Footnotes

1 Classified as a developed economy according to the World Economic Situation and Prospects 2024 report by the United Nations Department of Economic and Social Affairs (United Nations Department of Economic and Social Affairs, 2024, p. 135).

2 One of the infants in the Czech-exposed group was tested at the maternity ward of Havlíčkův Brod Hospital.

